# United States PRRSV 1-4-4 L1C.5 isolate demonstrates similar pathogenicity to a historic Chinese highly pathogenic PRRSV

**DOI:** 10.1101/2025.04.23.648216

**Authors:** Jayne E. Wiarda, Sarah J. Anderson, Hanjun Kim, Tyron Chang, Lauren Tidgren Hanson, Eraldo L. Zanella, Bailey Arruda, Meghan Wymore Brand, Samantha J. Hau, Jianqiang Zhang, Alexandra C. Buckley

**Affiliations:** Virus and Prion Research Unit, National Animal Disease Center, Agricultural Research Service, United States Department of Agriculture, Ames, Iowa, USA; Oak Ridge Institute of Science and Education, Agricultural Research Service Participation Program, Oak Ridge, Tennessee, USA; Department of Veterinary Diagnostic and Production Animal Medicine, College of Veterinary Medicine, Iowa State University, Ames, Iowa, USA

## Abstract

Porcine reproductive and respiratory syndrome virus (PRRSV) is a major economic and animal health burden on the United States swine industry due to morbidity- and mortality-associated losses affecting all stages of pig production. Currently, a large proportion of losses are attributed to a highly virulent PRRSV strain, PRRSV 1-4-4 L1C.5. To benchmark the virulence of PRRSV 1-4-4 L1C.5, a study was conducted to compare pathogenicity of this contemporary strain to a historical Chinese highly pathogenic PRRSV (HP-PRRSV) strain, JXwn06, that devastated the Chinese and other Asian swine industries since 2006, as well as a moderately virulent United States PRRSV strain, MN184, considered to be one of the most virulent PRRSV strains circulating in the United States in the early 2000s. Weaned pigs were inoculated with PRRSV strains L1C.5, JXwn06, MN184, or mock inoculum and necropsied at 2, 6, and 10 days post inoculation, or as needed due to severe disease. Clinical metrics, viral loads, cytokine concentrations, PRRSV-specific antibody concentrations, and pathology were compared between treatment groups to compare pathogenicity. Results indicate a high degree of similarity disease dynamics between L1C.5 and JXwn06 animals that diverged from MN184 and mock animals. Findings indicate L1C.5 and JXwn06 cause more severe morbidity and mortality in weaned pigs than MN184. Results may be applied to develop more effective strategies for mitigating PRRSV 1-4-4 L1C.5 outbreaks currently plaguing the United States swine industry.

## INTRODUCTION

Porcine reproductive and respiratory syndrome virus (PRRSV) causes one of the most economically important diseases affecting the swine industry. Clinical disease is characterized by abortions and other reproductive issues in breeding animals, as well as respiratory disease in all ages of growing pigs. PRRSV is an enveloped, positive-strand RNA virus in the genus *Betaarterivirus* that primarily targets macrophages for infection (Brinton et al., 2021). High mutation rate as well as recombination events lead to the consistent generation of new PRRSV strains, which can impede disease control measures, such as vaccination (Risser et al., 2021).

Recent data demonstrate growing economic losses attributed to PRRSV in the United States (U.S.), indicating PRRSV is becoming increasingly problematic to the U.S. swine industry. From 2006 to 2010, PRRSV cost the U.S. an estimated $664 million in annual losses (Holtkamp et al., 2013), and numbers have nearly doubled to an estimated $1.2 billion in annual losses between 2016 to 2020 based on preliminary reports (Farmer, 2024; O.H. Osemeke, 2024). Not only does PRRSV cause an economically devastating disease for the swine industry, but PRRSV also has an increasing negative impact on animal welfare. Contemporary PRRSV strains are reported to cause significantly higher morbidity and mortality than historic U.S. strains, which bodes poorly for future economic prosperity, food security, and animal health. In 2020, there were reports of high mortality events in U.S. finishing pigs that were attributed to a PRRSV strain with a 1-4-4 RFLP pattern that clustered in the L1C sublineage (specifically L1C.5) (Rawal et al., 2023; Yim-Im et al., 2023). The level of virulence for recently emerged PRRSV 1-4-4 L1C.5 strains was speculated to be comparable to highly pathogenic PRRSV (HP-PRRSV) strains that began circulating in China and Vietnam around 2006 and had a huge detrimental impact on the Asian swine industry (An et al., 2011; Tian et al., 2007), though such an *in vivo* comparison has not been made.

The objective of this study was to compare the pathogenicity of a recently emerged PRRSV 1-4-4 L1C.5 strain from the U.S. to historic examples of virulent PRRSV outbreaks identified in the early 2000s. Comparison strains included a well-documented, exceptionally virulent HP-PRRSV strain, JXwn06, that emerged in China in the early 2000s and devastated the Chinese and other Asian swine industries due to high morbidity and mortality events (Ruedas-Torres et al., 2021; Zhou et al., 2009; Zhou et al., 2008) and a moderate pathogenicity PRRSV strain, MN184, that circulated through the U.S. in the early 2000s and was considered to be a more virulent PRRSV strain compared to other strains circulating through the U.S. during that time (Brockmeier et al., 2012; Johnson et al., 2004; Ruedas-Torres et al., 2021). Due to peak respiratory disease caused by PRRSV being most commonly identified in pigs 4 to 10 weeks of age (Zimmerman et al., 2019), recently weaned pigs were used to evaluate metrics of pathogenicity for L1C.5, JXwn06, and MN184 PRRSV strains in this study. Overall, study results provide a greater understanding of the commonalities and divergences in disease dynamics caused by diverse PRRSV strains that will help guide future research efforts generating novel control and prevention measures against PRRSV.

## MATERIALS & METHODS

### Ethics statement

All animal work was performed following protocol ARS-24-1168 that was approved by the National Animal Disease Center (NADC) Institutional Animal Care and Use Committee. At scheduled necropsies or when an animal reached pre-determined humane euthanasia endpoints, animals were euthanized with intravenous administration of a barbiturate (Fatal Plus, Vortech Pharmaceuticals, Dearborn, MI) following the labeled dose.

### Inoculum preparation

Viral isolates used in this study were grown on MARC-145 cells cultured in minimum essential medium (MEM) supplemented with 10% fetal bovine serum at 37°C and 5% CO2. A plasmid was imported to the NADC containing a Chinese HP-PRRSV strain rJXwn06 (Guo et al., 2013). The virus (rJXwn06; GenBank accession EF641008.1) was rescued from the infectious clone and propagated on MARC-145 cells for five passages. Previously-performed pathogenicity studies using the rJXwn06 have confirmed high virulence comparable to the naturally-occurring JXwn06 Chinese HP-PRRSV strain (Guo et al., 2013), and we subsequently refer to the rescued JXwn06 virus (rJXwn06) as JXwn06. Two other U.S.-endemic PRRSV strains were provided by Dr. Zhang at Iowa State University. A PRRSV 1-4-4 L1C variant (L1C.5, USA/MN/01775/2021) that was passaged six times (GenBank accession OR634972.1) (Rawal et al., 2023), and a PRRSV 1-8-4 strain (MN184) that was passaged 13 times (GenBank accession EF488739.1) (Wang et al., 2008). All isolates were diluted in MEM to generate an inoculum titer of approximately 3 × 10^5^ TCID_50_/mL.

### Animal study design

Seventy-nine conventional, mixed breed, weaned pigs (∼3 weeks old) were purchased from a commercial farm in the U.S. Midwest and transported to the NADC in Ames, Iowa. Monthly health screening for the source farm was negative for PRRSV, *Mycoplasma hyopneumoniae*, and porcine epidemic diarrhea virus. Animals were then allocated into four treatment groups (n=20 for L1C.5, MN184, and mock; n=19 for JXwn06), considering balance of variables for weight and sex to minimize differences between treatment groups. A microchip (Bio-Thermo™, Destron Fearing, Dallas, TX) was inserted intramuscularly into each animal to measure body temperature. The JXwn06 group was housed in an ABLS-3Ag room due to its foreign origin, while the remaining groups (L1C.5, MN184, and mock) were housed in separate ABSL-2 rooms. All animals were provided ad-libitum feed and water. Animals were given 5 days of acclimation prior to inoculation. The JXwn06, L1C.5, and MN184 groups all received 2 mL of 3 × 10^5^ TCID_50_/mL viral solution intranasally with an atomization device (MAD Nasal™, Teleflex, Morrisville, NC) on 0 dpi. Similarly, the mock group received 2 mL of mock inoculum (MEM) intranasally.

Five animals from each treatment group were pre-assigned to necropsy timepoints at 2, 6, and 10 dpi, and necropsies were also performed on an as-needed basis if animals reached pre-determined humane euthanasia endpoints. The study ceased at 12 dpi due to complete loss of animals from L1C.5 and JXwn06 treatment groups. To reduce research animal usage, remaining animals in the MN184 and mock groups were transferred to another independent study at 12 dpi.

### Sample collection and processing

Body temperature was recorded daily using a microchip reader. Pigs were monitored for clinical signs daily and given clinical scores every other day. Pigs were given a respiratory score ranging from 0-3 (0= normal, 1= occasional cough/increased respiration, 2= coughing often/mild dyspnea, 3= productive cough/open-mouth breathing). Lethargy scores also ranged from 0-3 (0= normal, 1= moderate activity/normal abdominal fill, 2= slow moving/mildly gaunt, 3= laying down/no interest in food). Body weight was measured on 0, 6, and 10 dpi. Blood was collected via jugular venipuncture into serum separator tubes at 0, 2, 4, 6, and 10 dpi, centrifuged, and serum was aliquoted and stored at –80°C until further processing.

At necropsy, nasal swabs were collected by inserting a cotton swab into both nostrils and placing the swab in 2 mL viral transport media (Hank’s balanced salt solution with 2% fetal bovine serum, amphotericin B, and gentamicin). Bronchoalveolar lavage fluid (BALF) collection was performed after euthanasia by pipetting 50 mL of MEM into the trachea prior to the bifurcation to lavage both sides of the lungs and aspirating the fluid. To assess bacterial pneumonia, BALF was plated on brain heart infusion agar (Becton, Dickinson and Company, Franklin Lakes, NJ) and trypticase soy agar 5% sheep blood (Thermo Fisher Scientific, San Diego, CA). Plates were incubated at 37°C with 5% CO_2_ and examined at 24 and 48 hours. Bacterial growth was isolated by subculturing, and pure growth was submitted to the Diagnostic Bacteriology and Pathology Laboratory at the National Veterinary Services Laboratories in Ames, IA for MALDI-TOF identification. After collection and aliquoting, nasal swab and BALF samples were stored at –80°C until further processing.

At 10 dpi, tracheobronchial lymph node (TBLN) and lung sections from both cranial lobes and the left caudal lobe were collected for histopathology. Lung tissue with grossly identifiable lesions was collected when present. Histopathology tissues were fixed in 10% neutral-buffered formalin for ∼48 hours, followed by processing into formalin-fixed, paraffin-embedded (FFPE) tissue blocks. For RNA isolations to assess viral loads, TBLN, lung, thymus, olfactory bulb, nasal turbinate, and ethmoid were minced and placed into RNAlater Stabilization Solution (Thermo Fisher Scientific, San Diego, CA). Lung tissue with gross lesions was collected into RNAlater when identifiable; when gross lesions were not visible, lung tissue was collected from the left caudal lobe closest to the bronchial bifurcation. RNAlater tissues were stored at 4°C for ∼24 hours followed by transfer to –80°C storage until further processing.

### RNA isolation and reverse transcriptase quantitative PCR (RT-qPCR)

RNA was isolated from TBLN, lung, thymus, brain, nasal turbinate, and ethmoid stored in RNAlater. Samples were removed from –80°C storage and thawed to room temperature. For thymus, brain, and ethmoid, between 10-100 mg tissue was weighed and homogenized in Buffer RLT (QIAGEN, Santa Clarita, CA) containing 10 uL/mL beta-mercaptoethanol (350 uL/10 mg tissue) (Sigma Aldrich, Burlington, MA) using a tissue stator-rotor homogenizer (Cole-Palmer, Vernon Hills, IL) with hard tissue tips (Cole-Palmer, Vernon Hills, IL) to dissociate nasal turbinate or soft tissue tips (Cole-Palmer, Vernon Hills, IL) to dissociate thymus, olfactory bulb, and ethmoid. RLT homogenate was used as input for RNA isolations with the RNeasy Mini Kit with on-column DNase digestion according to manufacturer’s instructions (QIAGEN, Santa Clarita, CA). RNA was quantified using the RNA ScreenTape Analysis kit (Agilent, Santa Clara, CA) on a 4200 TapeStation System (Agilent, Santa Clara, CA). For TBLN and lung, 20-30mg tissue was homogenized in TRI Reagent (ThermoFisher San Diego, CA) using M-tubes (Miltenyi Biotech) on a GentleMACs OctoDissociator (Miltenyi Biotech), protocol RNA_01_01. Homogenate was used as input for RNA isolations with the MagMax-96 for Microarrays Total RNA Isolation kit following the spin procedure (Applied Biosystems, ThermoFisher San Diego, CA). RNA was quantified using the Bioanalyzer RNA Analysis kit (Agilent, Santa Clara, CA) on a 2100 Bioanalyzer system (Agilent, Santa Clara, CA). RNA was isolated from serum, nasal swab, and BALF using the MagMax-96 Viral RNA Isolation kit (Applied Biosystems, ThermoFisher San Diego, CA) following manufacturer instructions.

From isolated RNA, RT-qPCR was performed to quantify PRRSV viral copies as previously described (Miller et al., 2025). Briefly, the AgPath ID One Step RT-qPCR kit (Applied Biosystems, ThermoFisher San Diego, CA) was used with primers (PRRSV US-F: 5′- TCAGCTGTGCCAGATGCTGG, PRRSV US-R: 5′-AAATGGGGCTTCTCCGGGTTTT) and probe (5′-TCCCGGTCCCTTGCCTCTGGA) (Loving et al., 2007). Samples were run in duplicate on an Applied Biosystems 7500 Fast Real-Time PCR System (Applied Biosystems, ThermoFisher San Diego, CA). A standard curve of transcribed RNA was included on each plate to enable quantification of RNA copies. Input sample amounts for each well reaction included 800 ng RNA isolated from tissues and 8 uL isolated RNA from serum, BALF, and nasal swab collections.

### Cytokine detection

Cytokine concentrations for IFN-α, IFN-γ, TNF-α, IL-1β, IL-4, IL-6, IL-8, IL-10, and IL-12p40 were determined in serum and BALF samples using the ProcartaPlex™ Porcine Cytokine & Chemokine Panel 1, 9plex kit (ThermoFisher San Diego, CA). All serum and BALF samples were filtered using 96 well plates with a 40 μm mesh filter (MilliporeSigma, Burlington, MA) prior to the assay. 25 μL serum or BALF were diluted to target detectable assay ranges in universal assay buffer provided with the kit. The assay was carried out following the manufacturer’s protocol, and all samples were kept in the dark and incubated at 4°C overnight before measurement. Cytokine abundance was quantified using the Bio-Plex® MAGPIX™ Multiplex Reader (Biorad, Hercules, CA). Sample readings for IL-8 and IFN-γ were consistently over and under observable ranges, respectively, and were not able to be estimated confidently based on standard curves. Consequently, IL-8 and IFN-γ results were not included in further analysis.

Due to unobservable detection of IFN-γ in samples using the ProcartaPlex kit, IFN-γ was further measured in serum and BALF using a commercial enzyme-linked immunosorbent assay (ELISA) kit (Porcine IFN-γ; R&D Systems, Inc., Minneapolis, MN). However, the ELISA also showed no/minimal detectable IFN-γ levels in serum. Consequently, IFN-γ concentration are only provided for BALF and not serum for this work. Lack of IFN-γ detection may be attributable due to either a biological lack of the cytokine in samples or a technical issue related to signal detection interference in serum samples.

### PRRSV-specific antibody detection

A commercial ELISA kit (IDEXX PRRS X3 Ab Test, IDEXX Laboratories, Inc., Westbrook, ME) was used to detect PRRSV-specific antibody titers in serum and BALF samples. The assay is designed to detect antibodies in serum samples, so BALF samples were analyzed on an experimental basis. The assay was performed according to the manufacturer’s protocol. The plates were read at 650 nm using a SpectraMax 190 Microplate Reader (Molecular Devices, LLC, San Jose, CA) controlled by SoftMax Pro Software 7.0.2 (Molecular Devices, LLC). Optical density (OD) values were recorded, and ELISA results were expressed as sample-to-positive (S/P) OD ratios based on the manufacturer’s guidelines, with a positive result defined as an S/P ratio ≥ 0.4.

### Pathologic assessment and scoring

Following euthanasia, the pluck was removed, cleaned, and the dorsal and ventral surfaces were photographed. Based on previous animal challenge work with JXwn06 and an L1C.5 isolate (Brockmeier et al., 2017; Rawal et al., 2023), gross lung lesions were scored for comparison on 10 dpi. For gross lung lesion scores, the right cranial, right middle, right caudal, accessary, left caudal, caudal part of the left cranial, and cranial part of the left cranial lobe were each individually assigned a percentage of affected tissue surface area. A weighted score was then calculated to reflect volume percentage by lobe as previously described (Halbur et al., 1995).

### Statistical analysis

Data were compared between treatment groups within a single timepoint using the pairwise Wilcox test function, *pairwise.wilcox.test()*, available from the stats base package in R v4.2.1 (Team, 2022). Within each timepoint, all pairwise comparisons were made between treatment groups and corrected using the false discovery rate (FDR) method. Adjusted p-values <0.05 were considered significant. Significance was indicated in plots as *(<0.05), **(<0.005), or ***(<0.0005).

### Principal component analysis

Two subsets of data were utilized to perform principal component analysis (PCA) testing for data dimensionality reduction. The first dataset included all animals necropsied at 2, 6, or 10 dpi. The second dataset included only animals necropsied at 10 dpi that had increased longitudinal sampling performed. Parameters with detectable measurements for all samples and non-zero variability amongst samples within a dataset were identified and used to create principal components (PCs). The function, *prcomp()*, available from the stats base package in R v4.2.1 (Team, 2022) was used to calculate PCs. The percent contribution of each parameter to each PC was calculated by dividing the absolute value of each parameter’s rotation value by the sum of all parameter rotation values within each PC.

## RESULTS & DISCUSSION

### PRRSV strains JXwn06 and L1C.5 cause severe clinical disease and associated mortality

Humane endpoints were reached resulting in euthanasia of animals within the JXwn06 group between 7 to 10 dpi and in the L1C.5 group between 9 to 12 dpi, while no MN184 or mock animals were euthanized due to severe morbidity (**Figure 1A**). The study ceased at 12 dpi because of the loss of all animals from JXwn06 and L1C.5 groups due to a combination of morbidity-associated and scheduled (2, 6, 10 dpi) euthanasia.

**Figure 1.**
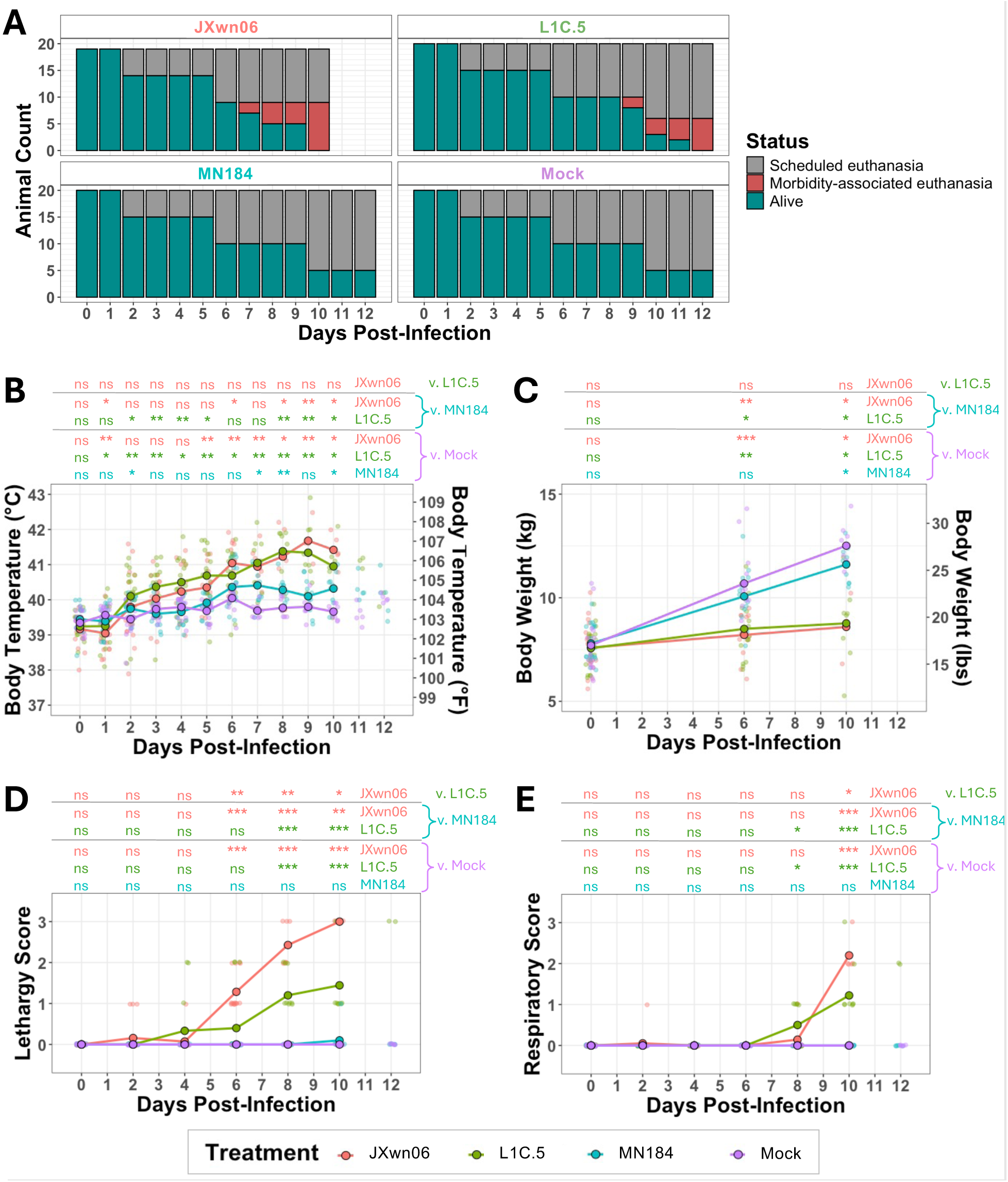
PRRSV strains L1C.5 and JXwn06 cause severe clinical morbidity and mortality in weaned pigs. **(A)** Counts (y-axes) for animals within each treatment group (individual panels) taken daily from 0 to 12 dpi (x-axes). Bar height represents the total number of animals in a treatment group, and bar color indicates the status of the animal. Data for the Jxwn06ic treatment group is not shown at 11 or 12 dpi due to all animals being euthanized by 10 dpi. **(B-E)** Measurements (y-axes) for body temperature **(B)**, body weight **(C)**, clinical scores for lethargy **(D)**, and clinical scores for respiratory signs **(E)** taken at indicated timepoints between 0 and 12 dpi (x-axes). Data were collected from all animals available at indicated timepoints. Mean values are shown as colored points and connecting lines for each treatment group at each timepoint. For data taken at 11 or 12 dpi, mean values are not shown due to no/low animal numbers in some treatment groups. Smaller, transparent points indicate individual animal data for each treatment group at each timepoint. Results for all statistical comparisons between treatment groups are shown as columns for each timepoint above each plot. All animals used in the study are shown for applicable timepoints and were used to calculate means and statistical values. Statistical comparisons of data were performed for all pairwise comparisons of treatment groups within each timepoint using the post-hoc, two-sided, unpaired Mann-Whitney U-test. Each timepoint was considered an individual dataset, and FDR correction of p-values was performed within each dataset (i.e. timepoint). Results based on FDR-corrected p-values are indicated as ns (p>=0.05), * (p<0.05), ** (p<0.005), or *** (p<0.0005). Abbreviations: dpi (days post-infection); FDR (false discovery rate); ns (not significant); PRRSV (porcine reproductive and respiratory syndrome virus); v. (versus)

To assess clinical disease following PRRSV challenge, body temperature, body weight, and clinical signs of lethargy and respiratory disease were recorded throughout the duration of the study (**Figure 1B-E**). All three PRRSV-inoculated groups had significantly elevated body temperatures compared to mock animals at various dpi. Additionally, JXwn06 and L1C.5 pigs had significantly higher post-infection body temperatures compared to MN184 animals (**Figure 1B**). Body weights were significantly lower in all three PRRSV-inoculated groups relative to mock animals by 10 dpi, and body weights for JXwn06 and L1C.5 pigs were also significantly lower relative to mock or MN184 animals at both 6 and 10 dpi (**Figure 1C**). Lethargy scores were significantly higher for JXwn06 animals starting at 6 dpi and L1C.5 animals starting at 8 dpi when compared to either mock or MN184 groups (**Figure 1D**). JXwn06 animals also had significantly higher lethargy scores than L1C.5 pigs beginning on 6 dpi (**Figure 1D**). Significantly higher respiratory scoring was recorded for L1C.5 pigs by 8 dpi and JXwn06 pigs at 10 dpi compared to both mock or MN184 animals (**Figure 1E**). Clinical scoring demonstrated MN184 animals had no disease-induced lethargy or respiratory distress. Signs of lethargy preceded respiratory signs in both LC1.5 and JXwn06 groups, and clinical signs developed earlier and were more severe in JXwn06 compared to L1C.5 animals.

The timeline for onset of clinical signs and animals reaching humane endpoints for euthanasia was expedited by ∼2 days for JXwn06 compared to L1C.5 animals, and onset of significantly higher scoring for clinical signs preceded morbidity-associated euthanasia of animals by one day for both treatment groups. Data demonstrate JXwn06 and L1C.5 are highly virulent PRRSV strains associated with severe morbidity that can cause mortalities in a relatively short time course of up to 12 days. Through 2024 and into 2025, L1C.5 isolates have been the predominate circulating strain detected in swine PRRSV diagnostic cases according to data from the Swine Disease Reporting System (Zeller MA et al., 2025). However, morbidity and mortality outcomes in the field may differ from outcomes observed in this study due to high variability in factors such as previous PRRSV exposure/vaccination, viral co-infections and/or secondary bacterial infections, antibiotic treatment, housing conditions, and environmental stressors that are either less controlled or less documented in a field setting.

### Higher viral loads are detected at systemic and local tissue sites in pigs challenged with PRRSV strains JXwn06 and L1C.5

Viral loads were assessed at systemic and local sites to compare tissue tropism and dissemination of PRRSV challenge strains. Quantification of viral RNA copies was completed in serum to detect viremia, in nasal swabs to detect potential viral shedding, in BALF, lung, nasal turbinate, ethmoid, and TBLN to indicate respiratory infection, and in thymus and olfactory bulb to indicate extra-respiratory spread (**Figure 2, 3**). PRRSV RNA was not detected in any mock-challenged animal samples.

**Figure 2.**
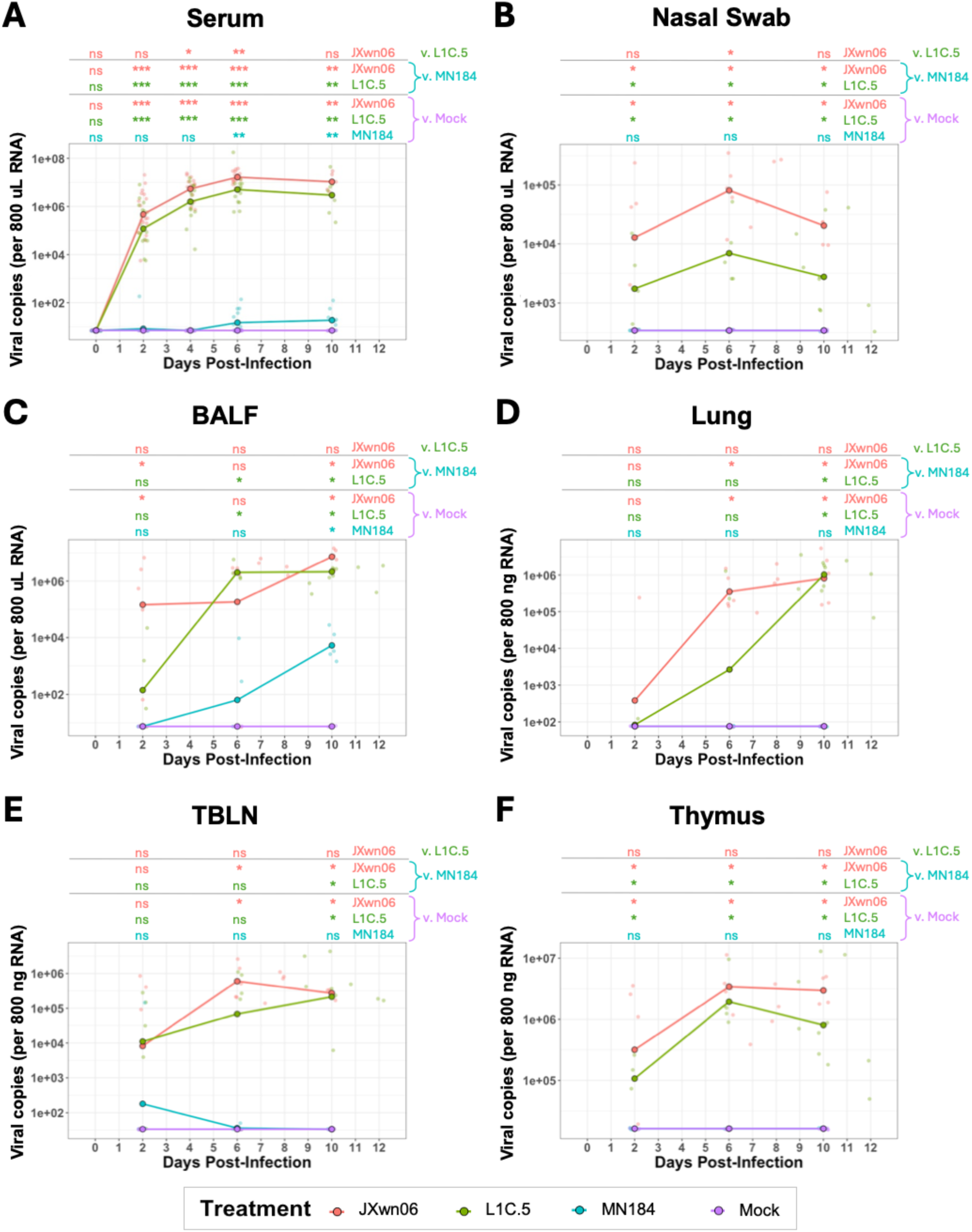
PRRSV strains L1C.5 and JXwn06 cause increased local and systemic viral loads. **(A-F)** Quantification of PRRSV RNA (y-axes) in serum **(A)**, nasal swabs **(B)**, BALF **(C)**, lung **(D)**, TBLN **(E)**, and thymus **(F)** taken at indicated timepoints between 0 and 12 dpi (x-axes). Serum data **(A)** were collected from all animals available at indicated timepoints. Nasal swab and tissue data **(A-F)** were collected from each animal at necropsy. Mean values are shown as colored points and connecting lines for each treatment group at each timepoint. For data taken outside of 0, 2, 4, 6, or 10 dpi for **(A)** or 2, 6, or 10 dpi in **(B-F)**, mean values are not shown due to no/low animal numbers in some treatment groups. Smaller, transparent points indicate individual animal data for each treatment group at each timepoint. Results for all statistical comparisons between treatment groups are shown as columns for each timepoint above each plot. All animals used in the study are shown for applicable timepoints and were used to calculate means and statistical values. Statistical comparisons of data were performed for all pairwise comparisons of treatment groups within each timepoint using the post-hoc, two-sided, unpaired Mann-Whitney U-test. Each timepoint was considered an individual dataset, and FDR correction of p-values was performed within each dataset (i.e. timepoint). Results based on FDR-corrected p-values are indicated as ns (p>=0.05), * (p<0.05), ** (p<0.005), or *** (p<0.0005). Abbreviations: BALF (bronchoalveolar lavage fluid), dpi (days post-infection); FDR (false discovery rate); ns (not significant); PRRSV (porcine reproductive and respiratory syndrome virus); TBLN (tracheobronchial lymph node), v. (versus)

PRRSV RNA was detected in serum samples of all three PRRSV treatment groups by 2 dpi. While serum viral copies for JXwn06 and L1C.5 groups were significantly higher than both mock and MN184 animals from 2 through 10 dpi, significantly higher values in MN184 compared to mock animals were not observed until 6 dpi but were also then sustained through 10 dpi (**Figure 2A**). The number of viral copies measured in serum was also significantly higher in JXwn06 relative to L1C.5 at 4 and 6 dpi (**Figure 2A**).

Viral copies were only detected in nasal swabs of JXwn06 and L1C.5 pigs starting at 2 dpi (**Figure 2B**). The highest average viral load was detected at 6 dpi for both JXwn06 and L1C.5 pigs, and viral copy numbers were significantly higher for JXwn06 compared to L1C.5 at this timepoint (**Figure 2B**).

Viral copy numbers in BALF, lung, and TBLN were significantly higher in JXwn06 and L1C.5 relative to both MN184 and mock groups at 10 dpi (**Figure 2C-E**). At 6 dpi, viral copies in lung and TBLN were also significantly higher for JXwn06 relative to both MN184 and mock animals (**Figure 2D-E**), and viral copies in BALF were significantly higher for L1C.5 animals relative to MN184 and mock (**Figure 2C**). Detection of viral copies in BALF but not lung of MN184 pigs suggests BALF may be a more reliable readout of PRRSV presence in lung, as detection may be severely skewed by the specific section of lung collected for analysis. Viral copies obtained from BALF, lung, or TBLN of JXwn06 versus L1C.5 did not show statistically significant differences at any timepoints.

Unlike respiratory tissues, thymus had significantly elevated viral copy numbers for JXwn06 and L1C.5 groups relative to both mock and MN184 groups as early as 2 dpi, and these significant elevations were maintained at 6 and 10 dpi (**Figure 2F**). Viral detection in thymus of MN184 pigs was minimal and did not significantly differ from mock animals at any timepoint (**Figure 2F**).

At 10 dpi, olfactory bulb, nasal turbinate, and ethmoid tissues were also sampled for PRRSV to investigate potential broadened tissue tropism to neurologic and nasal cavity sites. Similar levels of viral RNA were detected in the olfactory bulb of JXwn06 and L1C.5 pigs, and no viral RNA was detected in the olfactory bulb from MN184 or mock animals (**Figure 3**). Similarly, nasal turbinate and ethmoid demonstrated virus was present in these tissues for JXwn06 and L1C.5 but not MN184 pigs (**Figure 3**).

**Figure 3.**
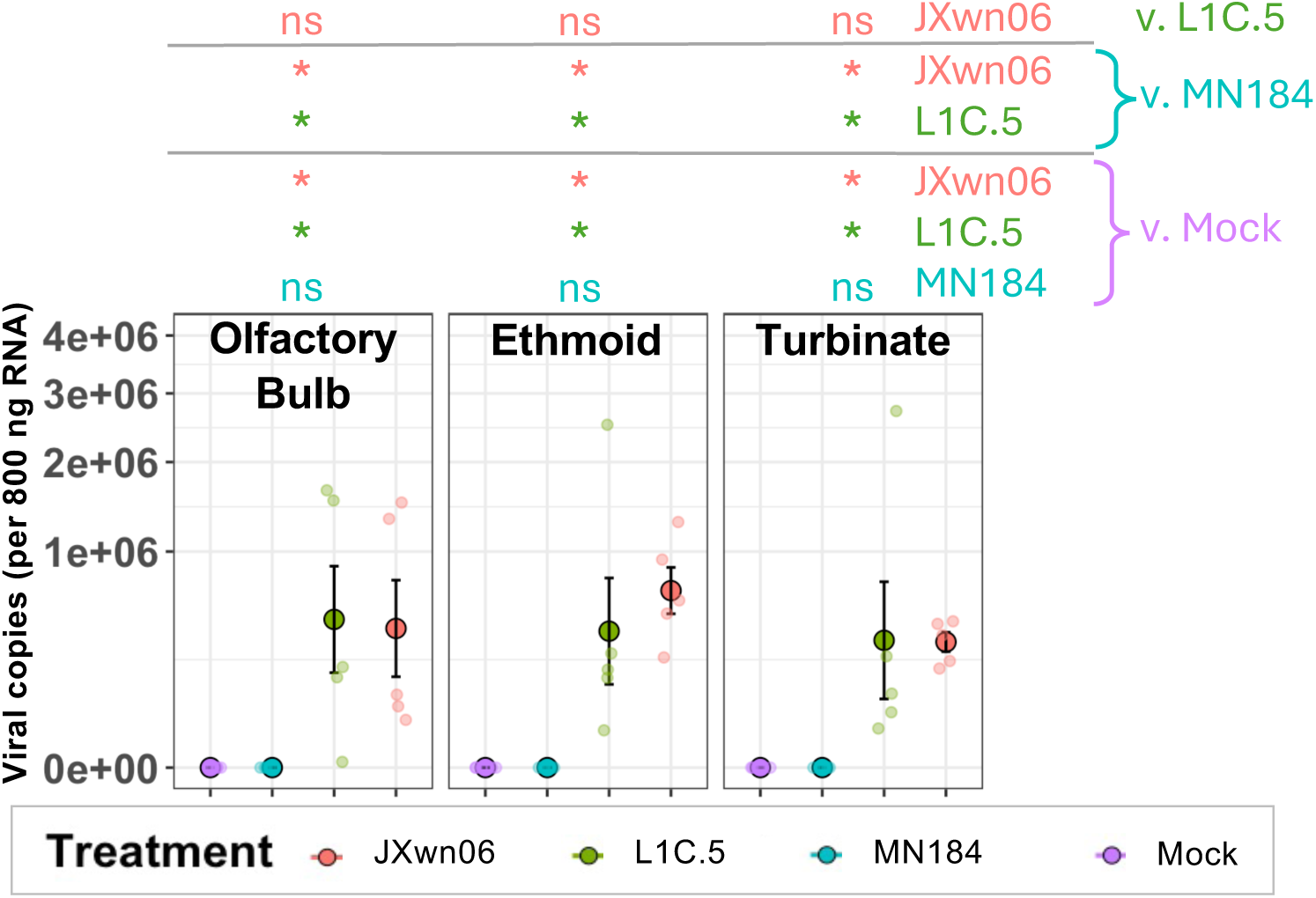
PRRSV strains L1C.5 and JXwn06 have broader nasal cavity and neurologic tissue tropism. Quantification of PRRSV RNA (y-axes) in olfactory bulb (left), ethmoid (center), and nasal turbinate (right) taken at 10 dpi from pigs of different treatment groups (x-axes). Mean values are shown as larger colored points for each treatment group within each tissue, and lines indicate standard errors. Smaller, transparent points indicate individual animal data for each treatment group in each tissue. Results for all statistical comparisons between treatment groups are shown as columns for each tissue above each plot. All animals euthanized at 10 dpi are shown and were used to calculate means, standard errors, and statistical values. Statistical comparisons of data were performed for all pairwise comparisons of treatment groups within each tissue using the post-hoc, two-sided, unpaired Mann-Whitney U-test. Each tissue was considered an individual dataset, and FDR correction of p-values was performed within each dataset (i.e. tissue). Results based on FDR-corrected p-values are indicated as ns (p>=0.05), * (p<0.05), ** (p<0.005), or *** (p<0.0005). Abbreviations: dpi (days post-infection); FDR (false discovery rate); ns (not significant); PRRSV (porcine reproductive and respiratory syndrome virus); v. (versus)

Results collectively suggest pigs challenged with JXwn06 and L1C.5 have higher viral loads detected systemically (serum), at respiratory sites (BALF, lung, TBLN, nasal swab, nasal turbinate, ethmoid), and at extra-respiratory immune (thymus) and neurologic (olfactory bulb) sites. Findings corroborate a previous study establishing HP-PRRSV strain JXwn06 has extended tissue tropism that may contribute to higher pathogenicity (Li et al., 2012) and expands upon initial findings of L1C.5 tissue tropism (Rawal et al., 2023). Extended tissue tropism may be a conserved hallmark of highly virulent PRRSV strains that could ultimately contribute to the exacerbated morbidity and mortality seen with these infections.

### PRRSV strains JXwn06 and L1C.5 cause heightened systemic and local cytokine responses

Cytokine concentrations were measured in serum to detect systemic immune responses (**Figure 4**) and in BALF to detect localized, respiratory immune responses (**Figure 5**) elicited by PRRSV infection. Overall, MN184 pigs showed some alterations to cytokine concentrations across the infection time course, but JXwn06 and L1C.5 animals showed extremely heightened levels of both systemic (**Figure 4**) and local (**Figure 5**) cytokines.

**Figure 4.**
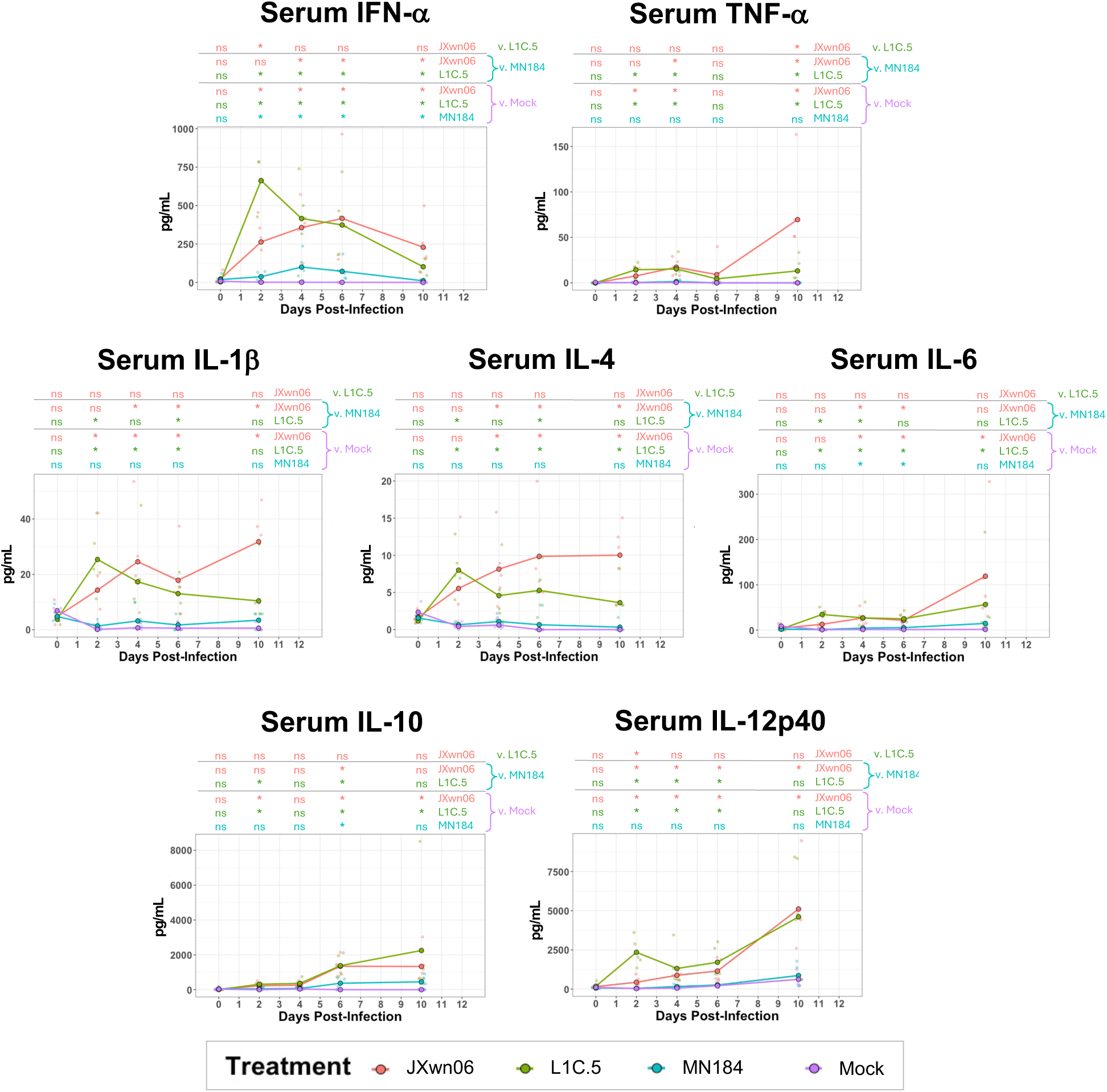
PRRSV strains L1C.5 and JXwn06 cause heightened systemic cytokine responses. Measurements (y-axes) for cytokine concentrations in serum taken at indicated timepoints of 0, 2, 4, 6, and 10 dpi (x-axes). Data were collected from all animals necropsied at 10 dpi (n=5/treatment group; n=20 total). Mean values are shown as colored points and connecting lines for each treatment group at each timepoint. Smaller, transparent points indicate individual animal data for each treatment group at each timepoint. Results for all statistical comparisons between treatment groups are shown as columns for each timepoint above each plot. All animals used in the study are shown for applicable timepoints and were used to calculate means and statistical values. Statistical comparisons of data were performed for all pairwise comparisons of treatment groups within each timepoint using the post-hoc, two-sided, unpaired Mann-Whitney U-test. Each timepoint was considered an individual dataset, and FDR correction of p-values was performed within each dataset (i.e. timepoint). Results based on FDR-corrected p-values are indicated as ns (p>=0.05), * (p<0.05), ** (p<0.005), or *** (p<0.0005). Abbreviations: dpi (days post-infection); FDR (false discovery rate); ns (not significant); PRRSV (porcine reproductive and respiratory syndrome virus); v. (versus)

**Figure 5.**
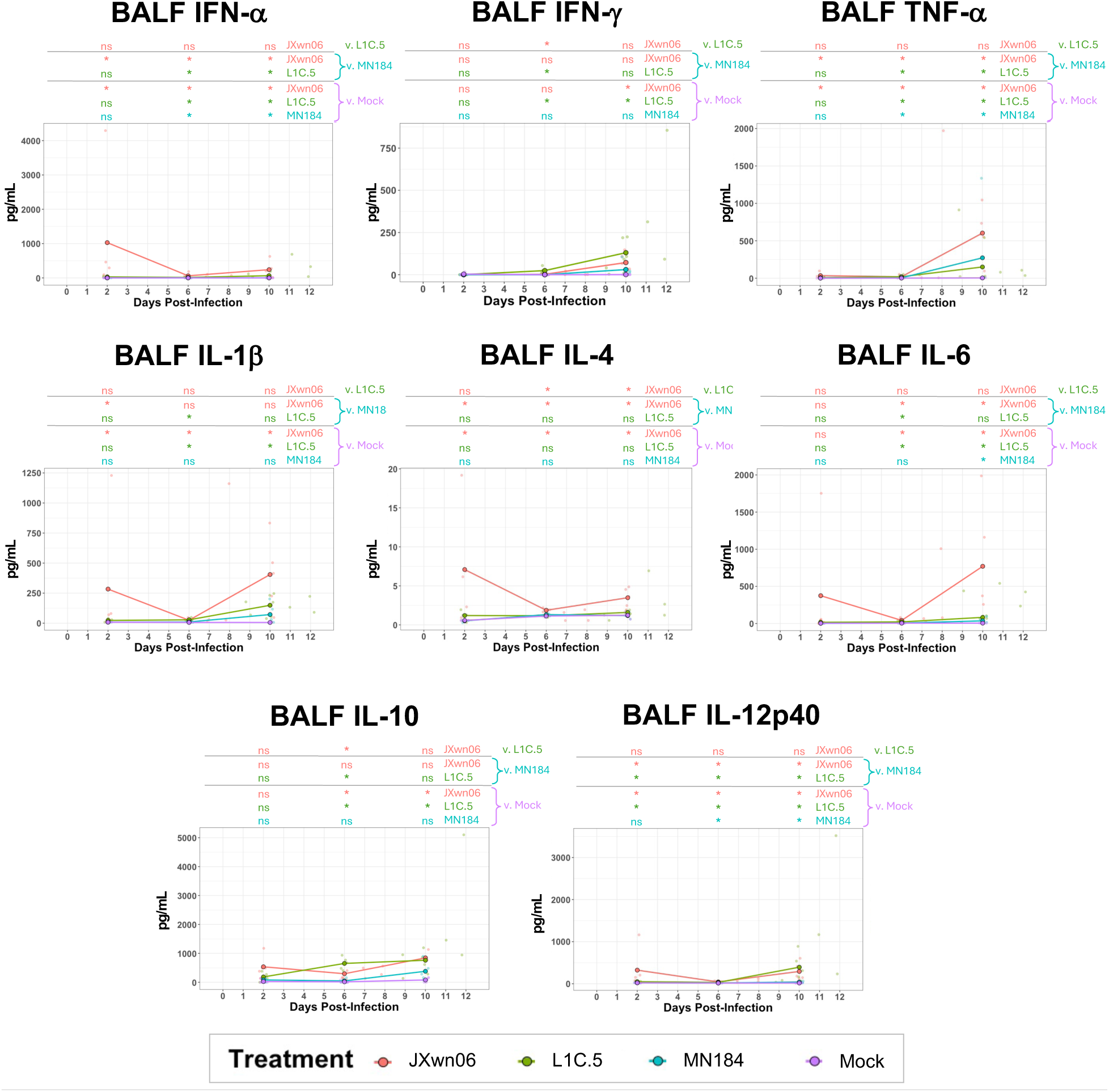
PRRSV strains L1C.5 and JXwn06 cause heightened localized, respiratory cytokine responses. Measurements (y-axes) for cytokine concentrations in BALF taken at indicated timepoints between 0 and 12 dpi (x-axes). Data were collected from each animal at necropsy. Mean values are shown as colored points and connecting lines for each treatment group at each timepoint. Smaller, transparent points indicate individual animal data for each treatment group at each timepoint. Results for all statistical comparisons between treatment groups are shown as columns for each timepoint above each plot. All animals used in the study are shown for applicable timepoints and were used to calculate means and statistical values. Statistical comparisons of data were performed for all pairwise comparisons of treatment groups within each timepoint using the post-hoc, two-sided, unpaired Mann-Whitney U-test. Each timepoint was considered an individual dataset, and FDR correction of p-values was performed within each dataset (i.e. timepoint). Results based on FDR-corrected p-values are indicated as ns (p>=0.05), * (p<0.05), ** (p<0.005), or *** (p<0.0005). Abbreviations: BALF (bronchoalveolar lavage fluid); dpi (days post-infection); FDR (false discovery rate); ns (not significant); PRRSV (porcine reproductive and respiratory syndrome virus); v. (versus)

In serum (**Figure 4**), cytokines in L1C.5 and/or JXwn06 pigs had significant elevations detected as early as 2 dpi that were sustained throughout the time course. In contrast, cytokine concentrations in MN184 pigs were only significantly higher relative to mock for IFN-α throughout the infection time course and for IL-6 and IL-10 at mid-infection timepoints (4, 6 dpi). Cytokine concentrations were relatively similar between L1C.5 and JXwn06 animals, with the exception of increased IFN-α and IL-12p40 in L1C.5 pigs at 2 dpi and elevated TNF-α in JXwn06 at 10 dpi.

In BALF (**Figure 5**), cytokine levels were also similar between L1C.5 and JXwn06 animals with the exception of IFN-ψ and IL-10 at 6 dpi and IL-4 at both 6 and 10 dpi. In general, the largest cytokine elevations relative to mock occurred at 10 dpi, including IFN-α, TNF-α, IL-6, and IL-12p40 concentrations being significantly higher than mock for all PRRSV treatment groups, while IFN-ψ, IL-1ý, and IL-10 concentrations were only significantly higher relative to mock for LIC.5 and JXwn06 animals.

Assessment of cytokine concentrations in serum and BALF collectively demonstrate both systemic and local immune responses were elicited by PRRSV infection, though cytokine responses were extremely heightened in animals inoculated with L1C.5 and JXwn06. Results coincide with previous reports of elevated pro-inflammatory and anti-inflammatory cytokine levels in serum and BALF of pigs infected with HP-PRRSV relative to lower virulence strains (Brockmeier et al., 2017; Guo et al., 2013). High elevation of both pro-inflammatory and anti-inflammatory cytokines in response to HP-PRRSV infections at 6 and 10 dpi may be indicative of widespread immune activation rather than a targeted, protective immune response.

Exacerbated immune activation may contribute to immunopathology-associated deterioration of an animal’s condition, as we observed increasing cytokine levels as disease progressed. Further analysis of the early immune response may enable us to identify immune processes predictive of disease resistance or reduced disease severity.

### PRRSV strains JXwn06 and L1C.5 cause heightened systemic and local virus-specific antibody responses

A PRRSV-specific humoral immune response was assessed by detecting PRRSV-specific antibodies in serum (**Figure 6A**) and BALF (**Figure 6B**) samples. PRRSV-specific antibodies were produced and began to circulate systemically between 6 to 10 dpi for all PRRSV treatment groups, as evidenced by the absence of detected antibody in any serum samples at 6 dpi and detection of similar antibody levels in MN184, L1C.5, and JXwn06 groups at 10 dpi (**Figure 6A**). In BALF, PRRSV-specific antibodies were similarly produced between 6 to 10dpi; however, antibodies were only detectable in L1C.5 and JXwn06 animals at 10 dpi (**Figure 6B**).

**Figure 6.**
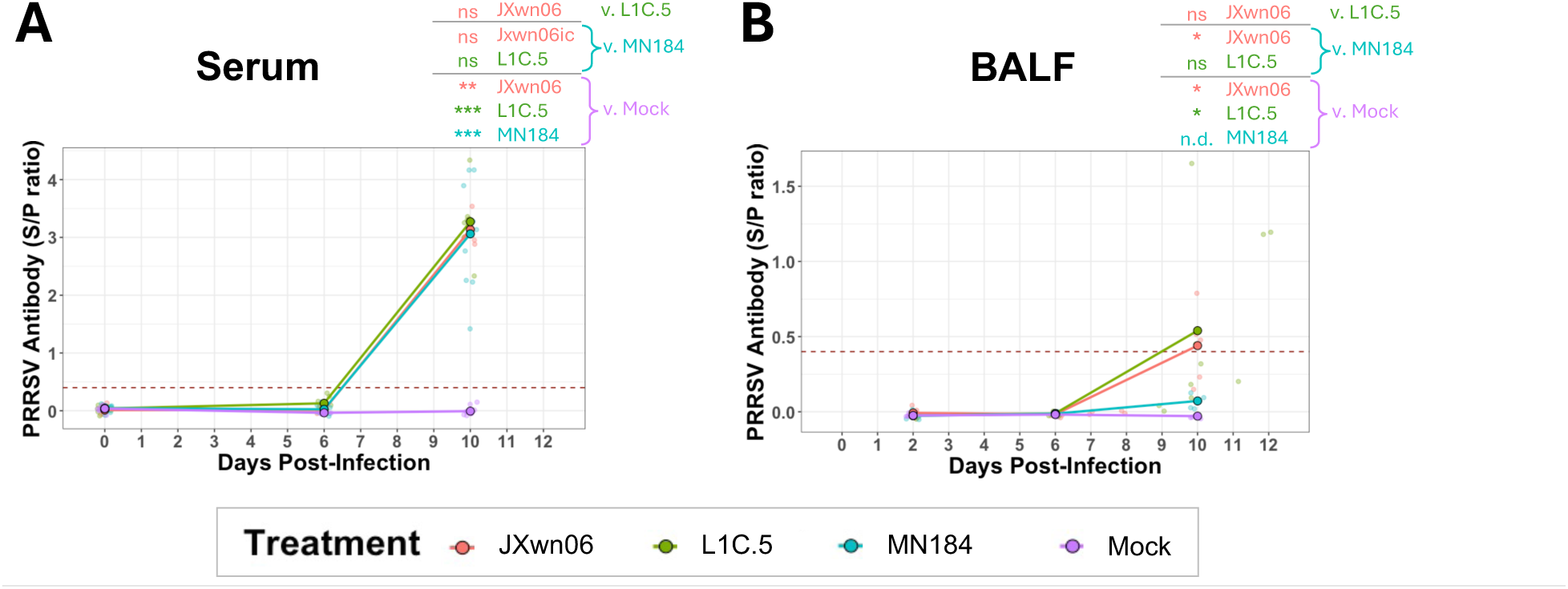
PRRSV strains L1C.5 and JXwn06 induce elevated levels of local and systemic anti-PRRSV antibodies. S/P values (y-axes) for PRRSV-specific antibody concentrations in serum **(A)** and BALF **(B)** taken at indicated timepoints between 0 and 12 dpi (x-axes). Serum data **(A)** were collected from all animals necropsied at 10 dpi (n=5/treatment group; n=20 total) at 0, 6, and 10 dpi. BALF data **(B)** were collected from each animal at necropsy. Mean values are shown as colored points and connecting lines for each treatment group at each timepoint. Smaller, transparent points indicate individual animal data for each treatment group at each timepoint. Results for all statistical comparisons between treatment groups are shown as columns for each timepoint above each plot. All animals used in the study are shown for applicable timepoints and were used to calculate means and statistical values. Statistical comparisons of data were performed for all pairwise comparisons of treatment groups within each timepoint using the post-hoc, two-sided, unpaired Mann-Whitney U-test. Each timepoint was considered an individual dataset, and FDR correction of p-values was performed within each dataset (i.e. timepoint). Statistical analysis was not performed if no values were above the level of detection confidence (S/P ratio > 0.4) within a timepoint. Results based on FDR-corrected p-values are indicated as ns (p>=0.05), * (p<0.05), ** (p<0.005), or *** (p<0.0005). Abbreviations: BALF (bronchoalveolar lavage fluid); dpi (days post-infection); FDR (false discovery rate); n.d. (not determined); ns (not significant); PRRSV (porcine reproductive and respiratory syndrome virus); v. (versus)

Results indicated successful generation of PRRSV-specific antibodies between 6 to 10 dpi, with all PRRSV-infected pigs no matter the virulence of the isolate generating similar levels of antibodies in serum by 10 dpi. Only pigs challenged with L1C.5 and JXwn06 had detectable PRRSV-specific antibody in BALF, suggesting distinctions in local humoral immunity could be dependent on PRRSV pathogenicity. Local humoral immunity producing PRRSV-specific antibody in L1C.5 and JXwn06 animals may be influenced by increased lung vascular permeability and/or elevated antibody production by lung-associated B cells (Mulupuri et al., 2008; Sun et al., 2022). Higher PRRSV load (**Figure 2C-D**) and heightened cytokine activity (**Figure 5**) in the lung environment may also act as stimulatory signals to promote lung-localized antibody production. However, changes in the pulmonary environment may instead be linked to increased lung immunopathology caused by higher pathogenicity PRRSV infections and may not necessarily correlate to increased antibody protection in the localized lung environment.

### Severe respiratory pathology observed with PRRSV strains JXwn06 and L1C.5

At 10 dpi necropsies, scoring was completed on lung lobes to assess gross pathology. The percent of lung surface area containing gross lesions was estimated (**Figure 7A**), a weighted average was calculated (**Figure 7B**), and representative images are shown in **Figure 7C**. Percentages of gross pathology for each lung lobe (**Figure 7A**) and weighted averages (**Figure 7B**) demonstrated similar results: a larger percentage of lung surface area contained gross lesions in L1C.5 and JXwn06 lungs. Though MN184 gross pathology was not significantly greater than mock, more non-zero gradings were observed in MN184 animals, and small group sizes likely contributed to low statistical power yielding a non-significant p-value of 0.1 when comparing weighted averages of MN184 and mock animals in **Figure 7B**.

**Figure 7.**
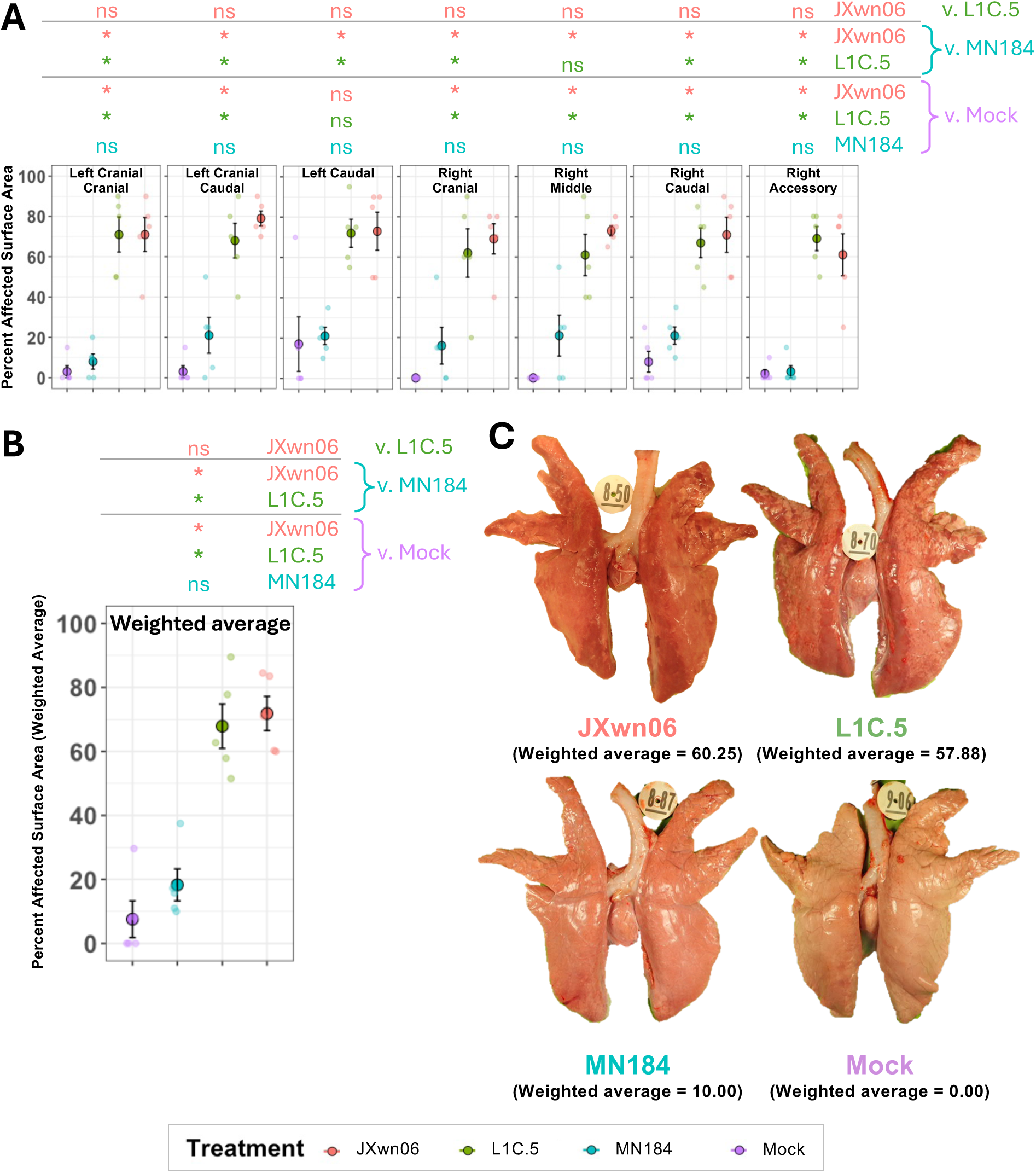
PRRSV strains L1C.5 and JXwn06 cause severe gross lung lesions at 10 dpi. **(A)** Quantification of percentage of lung surface area grossly exhibiting signs of pathology (y-axes) in individual lung lobes (**A**) or as a weighted average (**B**) taken at 10 dpi from pigs of different treatment groups (x-axes). Mean values are shown as larger colored points for each treatment group within each lung lobe panel, and lines indicate standard errors. Smaller, transparent points indicate individual animal data for each treatment group in each tissue. In **(C)**, representative images of gross lung pathology are shown for one animal of each treatment group. Within each treatment, pigs with the highest or lowest weighted average gross pathology scores were not selected as representative images. Results for all statistical comparisons between treatment groups are shown as columns for each tissue above each plot. All animals euthanized at 10 dpi (n=5/treatment group; n=20 total) are shown and were used to calculate means, standard errors, and statistical values. Statistical comparisons of data were performed for all pairwise comparisons of treatment groups within each tissue using the post-hoc, two-sided, unpaired Mann-Whitney U-test. Each tissue was considered an individual dataset, and FDR correction of p-values was performed within each dataset (i.e. tissue). Results based on FDR-corrected p-values are indicated as ns (p>=0.05), * (p<0.05), ** (p<0.005), or *** (p<0.0005). Abbreviations: dpi (days post-infection); FDR (false discovery rate); ns (not significant); PRRSV (porcine reproductive and respiratory syndrome virus); v. (versus)

We identified plum-colored consolidation in pigs from both the JXwn06 (n=3) and L1C.5 (n=2) groups consistent with bacterial bronchopneumia on 10 dpi. To enumerate bacterial loads, bacterial plating was performed on necropsy BALF samples. Although bacterial members causing porcine respiratory disease (*Bordetella bronchiseptica* and *Streptococcus suis*) were detected in several BALF samples across challenge groups and time points, bacterial plating results did not indicate a strain-specific increase in bacterial isolation (**Table 1**).

**Table 1.**
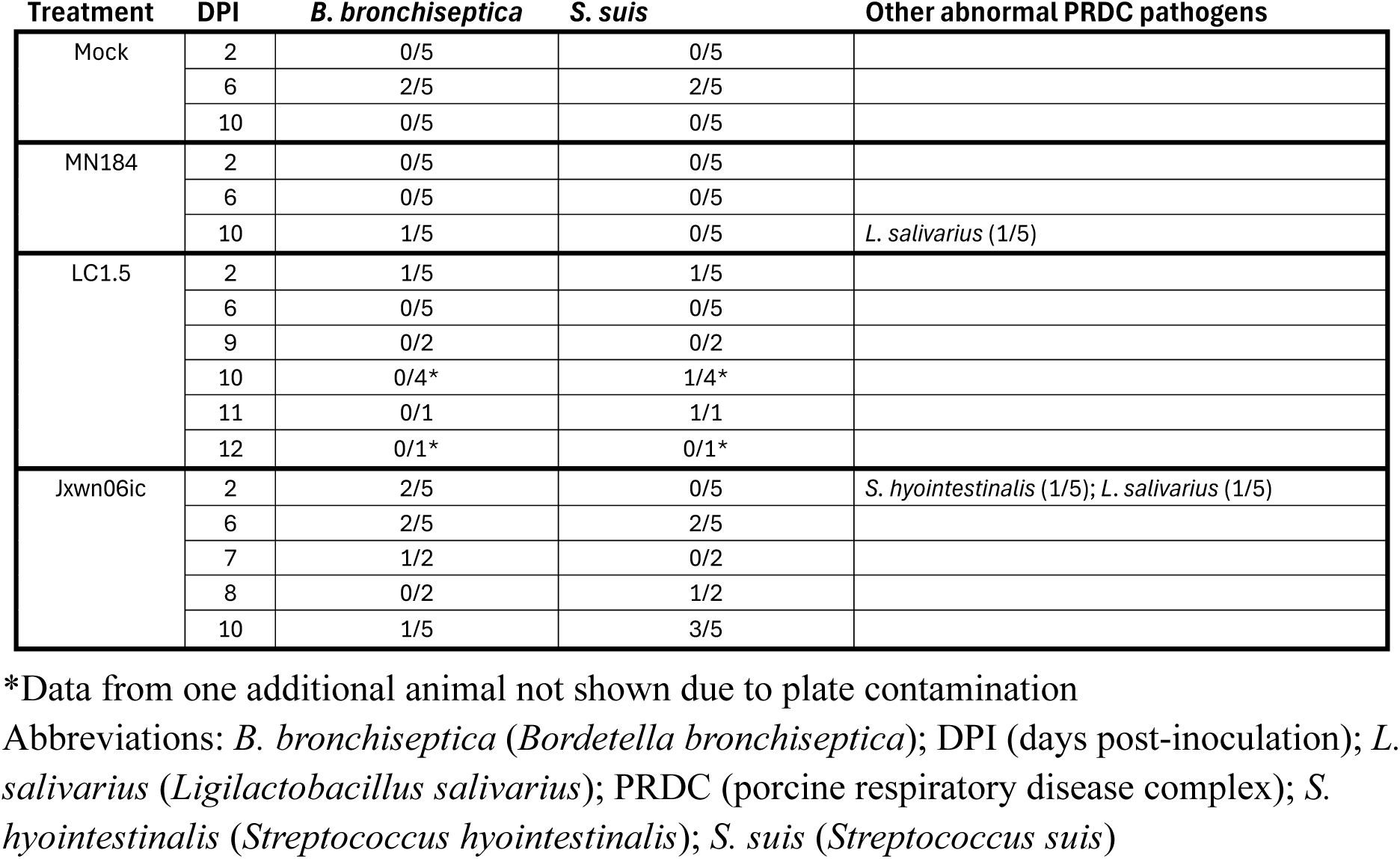
Bacterial plating of BALF samples.

PRRSV-associated lung pathology was greatest in pigs inoculated with L1C.5 and JXwn06. This is consistent with prior work demonstrating severe lung pathology in animals challenged with both L1C.5 (Rawal et al., 2023) and JXwn06 (Guo et al., 2013); however, there has been no direct comparisons of these strains. Additionally, prior work has shown PRRSV pathology may contribute to or be exacerbated by secondary bacterial infections (Brockmeier et al., 2017). In our study, gross lesion scoring evaluated the percent of lung affected and did not differentiate between viral interstitial pneumonia and bacterial pneumonia. While PRRSV is a major contributor to development of complicated porcine respiratory disease complex infections, it is difficult to recapitulate these complex infections in an experimental setting. Although we did not see a strong correlation between more severe PRRSV disease and development of secondary bacterial infections in this study, the impact of PRRSV-bacterial interactions may be different in a field setting where other stressors contribute to infection.

### A multi-parameter benchmark of virulence for contemporary PRRSV L1C.5 relative to historic HP-PRRSV strain JXwn06

To determine the overall relatedness of samples based on cumulative parameter contributions measured in preceding results, a multidimensional reduction analysis of data was undertaken. A dataset of samples with end-state readouts for pigs euthanized at 2, 6, and 10 dpi was created (**Figure 8**), and a second dataset included time-series data taken more comprehensively from pigs necropsied at 10 dpi (**Figure 9**).

**Figure 8.**
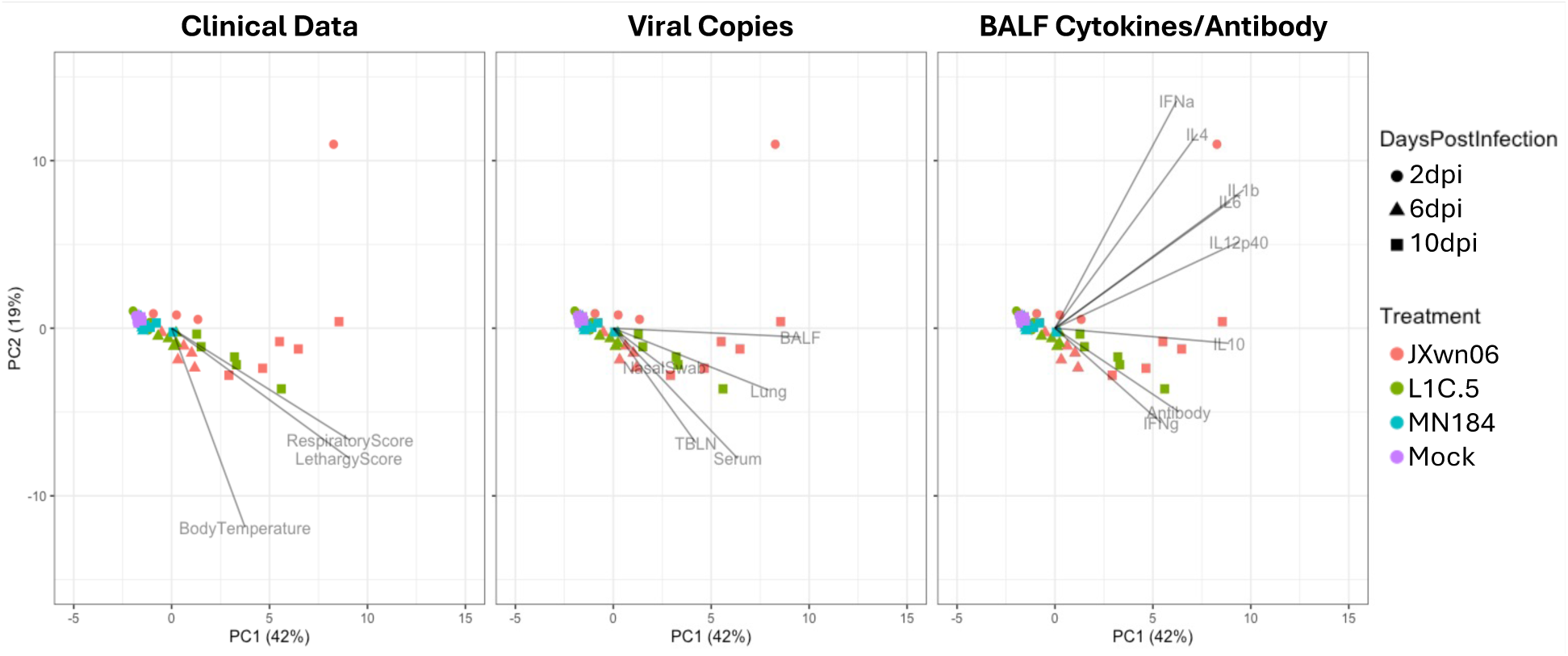
PRRSV strains L1C.5 and JXwn06 causes temporally progressive divergence from mock and MN184 samples across acute infection time course. Dimensional reduction (PCA analysis) of data collected from all animals necropsied at 2, 6, and 10 dpi. Points represent individual animals, where point color indicates treatment, and point shape indicates necropsy timepoint. Individual panels show contribution of PCs broken into general categories. The direction of a PC indicates positive correlation to samples diverging from the (0,0) coordinate in a similar direction. Longer lines indicate greater contribution of a component to the PCs and directions shown. Points for 60 animals are shown, including 15 animals from each treatment group, with 5 of each treatment group necropsied at each timepoint, and data from all 60 animals were used to calculate PCs. Abbreviations: BALF (bronchoalveolar lavage fluid); dpi (days post-infection); PCA (principal component analysis); PC (principal component); PRRSV (porcine reproductive and respiratory syndrome virus); TBLN (tracheobronchial lymph node)

**Figure 9.**
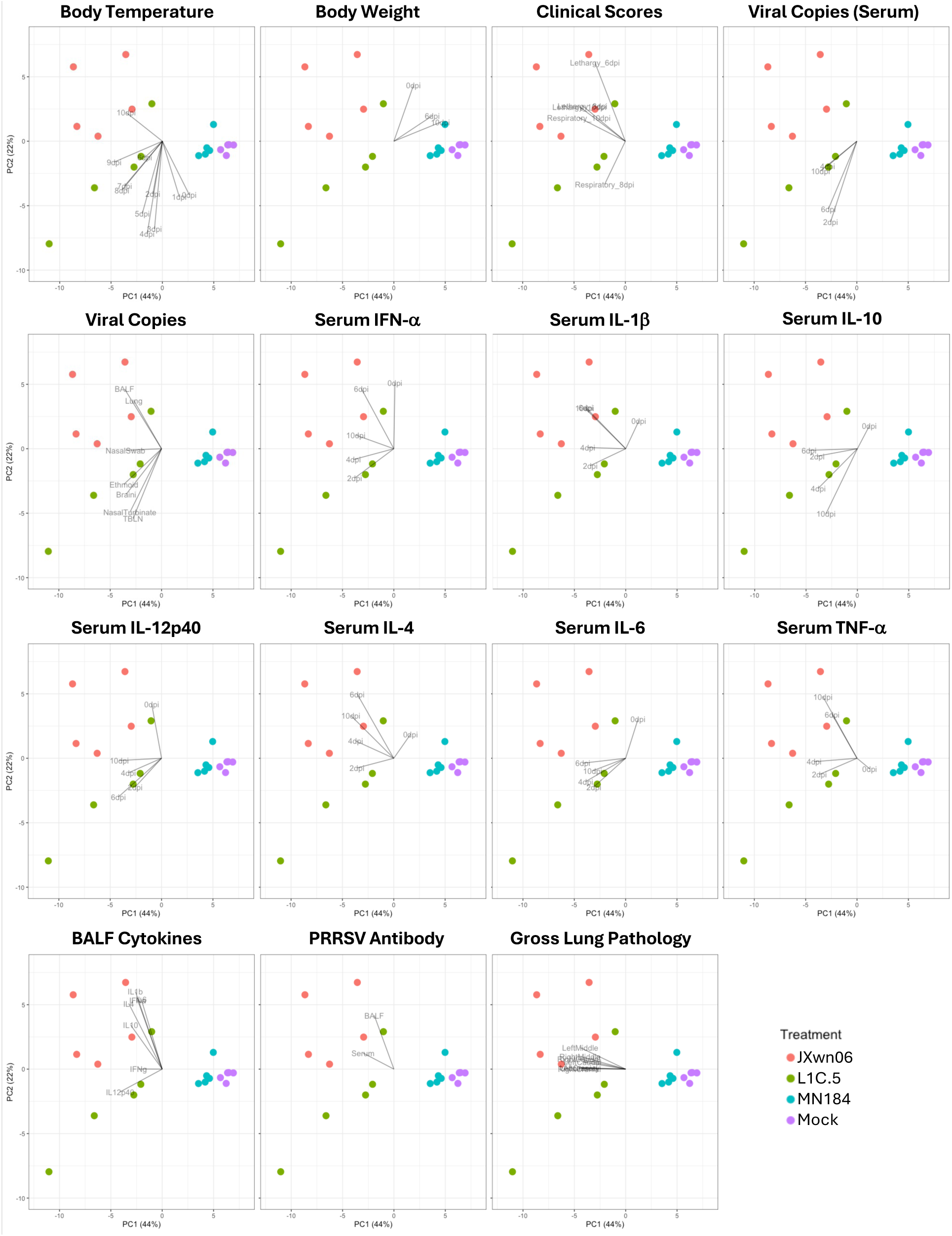
PRRSV strains L1C.5 and JXwn06 causes severe disease based on temporal and terminal readouts of host response. Dimensional reduction (PCA analysis) of data collected from pigs necropsied at 10 dpi across different treatment groups. Points represent individual animals, where point color indicates treatment. Individual panels show contribution of PCs broken into general categories. The direction of a PC indicates positive correlation to samples diverging from the (0,0) coordinate in a similar direction. Longer lines indicate greater contribution of a component to the PCs and directions shown. All animals euthanized at 10 dpi (n=5/treatment group; n=20 total) are shown and were used to calculate PCs. Abbreviations: BALF (bronchoalveolar lavage fluid); dpi (days post-infection); PRRSV (porcine reproductive and respiratory syndrome virus); PCA (principal component analysis); PC (principal component); TBLN (tracheobronchial lymph node)

PCA analysis of end-state readouts for pigs euthanized at 2, 6, and 10 dpi (**Figure 8**) indicated JXwn06 and L1C.5 samples progressively diverged from mock and MN184 samples as post-inoculation time increased. MN184 samples, regardless of dpi, remained more closely associated with mock samples, indicating a lower degree of divergence caused by MN184 infection relative to L1C.5 or JXwn06. Temporally-progressive divergence of JXwn06 and L1C.5 samples from mock and MN184 groups was mainly associated with PC1, with 6 and 10 dpi samples most largely diverging from mock samples. Clinical readouts, viral loads, and BALF cytokine/antibody readouts also showed a strong positive correlation with divergence of later-timepoint L1C.5/JXwn06 samples along PC1. One particular sample collected from a JXwn06 animal at 2 dpi had extremely high cytokine concentrations detected in BALF, causing it to be an outlier.

Analysis of time series data from 10 dpi-euthanized pigs (**Figure 9**) also indicated a large difference between L1C.5/JXwn06 and mock/MN184 samples, with mock and MN184 samples having tighter clustering and being located closer to one another, indicating a smaller degree of variation. In contrast, L1C.5/JXwn06 samples diverged from mock samples primarily along PC1 and did not have as tight of clustering amongst datapoints of a treatment group, indicating more within-group variability. Later dpi body temperatures, clinical scores, viral copies, later dpi cytokine concentrations, antibody concentrations, and gross lung pathology positively correlated with L1C.5 and JXwn06 samples along PC1, while early dpi temperature data, body weight, and 0 dpi cytokine concentrations had negative correlations.

JXwn06 and L1C.5 samples also exhibited notable divergence from each other along PC2, indicating divergence between the two higher pathogenicity PRRSV groups, though PC2 accounted for a smaller degree of variability (22%) compared to PC1 (44%). Along PC2, most clinical scores had a stronger positive correlation with JXwn06 samples, as did BALF/lung viral copies, several later dpi serum cytokines (IFN-α, IL-1ý, IL-4, TNF-α), most BALF cytokines, and PRRSV antibody. Alternatively, parameters including later dpi body temperatures, serum viral copies, viral copies in olfactory bulb/nasal turbinate/ethmoid/TBLN, and later dpi serum cytokines (IL-10, IL-12p40, IL-6) tended to have stronger positive correlations with L1C.5 samples.

Collectively, multi-parameter analyses indicated a strong divergence of JXwn06 and L1C.5 that suggest recently emerging United States PRRSV strain L1C.5 causes highly similar disease dynamics compared to early 2000s Chinese HP-PRRSV strain JXwn06, though some slight differences were noted. MN184, once considered one of the more virulent PRRSV strains circulating in the U.S. during the early 2000s, did cause disease but was much less virulent than L1C.5, demonstrating a trend towards emergence of more pathogenic PRRSV strains in the U.S. over the past two-and-a-half decades. Results establish a historic benchmark of pathogenicity for PRRSV L1C.5 that suggest similar disease severity and dynamics to Chinese HP-PRRSV strain JXwn06, which caused detrimental disease outbreaks in early 2000s China.

## DATA AVAILABILITY

Source data and computational code will be made available upon final publication.

## ACKNOWLEDGEMENTS

We thank the following for their valuable contributions to the work: (1) Colin Stoy, Andrew Von Weber, and Deborah Adolphson for technical assistance; (2) Justin Miller, Alyssa Bergeson, Randy Leon, Cam Nelson, Jason Huegel, Dr. Jean Kaptur, Dr. Rebecca Cox, Caitlyn Ehrlich, Jonathan Gardner, Tiffany Williams, Emma Hay, Emma Pratt, and Jared Peterson for animal care; (3) Adrienne Shircliff, Judith Stasko, and Sami Notebloom for histology expertise; (4) Denise Chapman for animal procurement; (5) the Animal Resources Unit and Institutional Animal Care and Use Committee at the National Animal Disease Center for oversight/approval of animal work.

## AUTHOR CONTRIBUTIONS

**Jayne E. Wiarda:** conceptualization, methodology, investigation, formal analysis, resources, data curation, visualization, supervision, project administration, writing – original draft, writing – review and editing. **Sarah J. Anderson:** methodology, investigation, formal analysis, writing – review and editing. **Hanjun Kim:** methodology, investigation, formal analysis, writing – review and editing. **Tyron Chang:** methodology, investigation, formal analysis, writing – review and editing. **Lauren Tidgren Hanson:** investigation, writing – review and editing. **Eraldo L. Zanella:** investigation, writing – review and editing. **Bailey Arruda:** methodology, investigation, formal analysis, writing – review and editing. **Meghan Wymore Brand:** methodology, investigation, formal analysis, resources, supervision, writing – review and editing. **Samantha J. Hau:** methodology, investigation, formal analysis, resources, supervision, writing – review and editing. **Jianqiang Zhang:** methodology, resources, writing – review and editing. **Alexandra C. Buckley:** conceptualization, methodology, investigation, formal analysis, resources, supervision, project administration, writing – original draft, writing – review and editing.

## CONFLICT OF INTEEST STATEMENT

Authors declare no conflict of interest.

## FUNDING

This work was funded by United States Department of Agriculture (USDA) Agricultural Research Service (ARS) CRIS project #5030-32000-230-000-D. This work was also funded by an appointment to the ARS Research Participation Program administered by the Oak Ridge Institute for Science and Education (ORISE) through an interagency agreement between the U.S. Department of Energy (DOE) and the USDA. ORISE is managed by Oak Ridge Associated Universities (ORAU) under DOE contract number DE-SC0014664. All opinions in the paper are the authors’ and do not necessarily reflect the policies and views of USDA, ARS, DOE, or ORAU/ORISE. Mention of trade names or products is for informational purposes only and does not imply endorsement by the USDA. USDA is an equal opportunity employer and provider.

## Notes

### Competing Interest Statement

The authors have declared no competing interest.

